# Factor Structure and Heritability of Obsessive-Compulsive Traits in Children and Adolescents in the General Population

**DOI:** 10.1101/246520

**Authors:** C.L. Burton, L. S. Park, E. C. Corfield, N. Forget-Dubois, A. Dupuis, V. M. Sinopoli, J. Shan, T. Goodale, S-M Shaheen, J. Crosbie, R.J. Schachar, P.D Arnold

## Abstract

**Background:** Obsessive-compulsive disorder (OCD) is a heritable childhood-onset psychiatric disorder that may represent the extreme of obsessive-compulsive (OC) traits that are widespread in the general population. We studied the factor structure and heritability of the Toronto Obsessive Compulsive Scale (TOCS), a new measure designed to assess traits associated with OCD in children and adolescents. We also examined the degree to which genetic effects are unique and shared between dimensions.

**Methods:** OC traits were measured using the TOCS in 16,718 children and adolescents (6 to 18 years) at a local science museum. Factor analysis was conducted to identify OC trait dimensions. Univariate and multivariate twin modeling was performed to estimate the heritability of OC trait dimensions in a subset of twins (220 pairs).

**Results:** Six OC dimensions were identified: Cleaning/Contamination, Hoarding, Rumination, Superstition, Counting/Checking, and Symmetry/Ordering. The TOCS total score (74%) and OC trait dimensions were heritable (30-77%). Hoarding was phenotypically distinct but shared genetic effects with other OC dimensions. Most of the genetic effects were shared between dimensions while unique environment accounted for the majority of dimension-specific variance, except for hoarding which had considerable unique genetic factors. A latent trait did not account for the shared variance between dimensions.

**Conclusions:** OC traits and individual OC dimensions were heritable, although the degree of shared and dimension-specific etiological factors varied by dimension. The TOCS is useful for genetic research of OC traits and OC dimensions should be examined individually and together along with total trait scores to characterize OC genetic architecture.

## INTRODUCTION

The role of genetics in the etiology of obsessive-compulsive disorder (OCD) is well-established (see Pauls et al., 2014 for a review). However, there are few replicated genetic risk variants for OCD. Gene discovery has been hampered by relatively small sample sizes, a deficiency which international consortia are working to overcome (International Obsessive Compulsive Disorder Foundation Genetics and Studies, 2017).

Despite progress, exclusive reliance on OCD diagnosis and clinic samples may impede progress. Diagnoses are useful in clinical practice, but could obscure phenotypic and genetic heterogeneity in study samples, hide variation in symptom severity among affected individuals, and miss sub-threshold OCD cases (Plomin et al., 2009). Clinical samples are slow and expensive to collect. One alternative is using quantitative OC trait measures that would assess the full range of OC traits (e.g., from extreme difficulty discarding useless objects to being able to easily discard useless objects) to capture all the variance in these behaviours. A quantitative trait-based measure could boost power for genetic studies (Plomin et al., 2009, van der Sluis et al., 2013), especially in general population samples.

Quantitative scores can be derived from traditional OCD scales, but most are symptom-based. Symptom counts typically generate J-shaped distributions that are suboptimal for quantitative analyses and are particularly problematic if measured in general population samples given most participants will have few or no OCD symptoms.

Another obstacle to genetic research in OCD is variability in the presentation of the disorder which suggests etiological heterogeneity. Phenotypic heterogeneity evident in factor analyses demonstrates that OCD symptoms generally cluster into four dimensions: symmetry, forbidden thoughts/checking, cleaning, and hoarding (Bloch et al., 2008, Stewart et al., 2008). Hoarding is often considered a distinct dimension and is classified as its own disorder in the most recent edition of the Diagnostic and Statistical Manual of Mental Disorders (DSM-5; American Psychiatric Association, 2013). In adults, twin studies indicate that phenotypic heterogeneity reflects etiological heterogeneity. Each dimension is heritable with unique and shared genetic influences contributing to their etiology (Katerberg et al., 2010, Iervolino et al., 2011, van Grootheest et al., 2007a). In children and adolescents, the heritability of obsessive-compulsive (OC) traits and dimensions is unclear (Moore et al., 2010).

Understanding the degree of shared and unique genetic influences on OC dimensions is critical to designing genetic studies to uncover the genetics of a heterogeneous trait. If OC dimensions are highly correlated with mostly shared genetic influences or if their shared variance is captured by a latent trait (e.g., global OC traits), than studies should focus on the latent trait. However, if OC dimensions are influenced independently by shared and unique genetic factors then studies should focus on both the overall trait and the dimensions. The best fitting model is unclear in adults (Iervolino et al., 2011, van Grootheest et al., 2008b) and has not been addressed in youth.

We developed the Toronto Obsessive Compulsive Scale (TOCS; Park et al., 2016) to measure the full range of OC traits in children and adolescents. To understand the utility of the TOCS for genetic research and explore the genetic influences on OC dimensions in children, we asked if the TOCS captured dimensions similar to traditional OCD measures, if these dimensions were heritable as well as co-heritable, and if these dimensions had shared and unique etiological factors that influenced each dimension independently or through a latent trait. To answer these questions, we conducted a factor analysis to identify OC dimensions in a population-based sample (n=16,718) and then we fit ACE models to the twins (n=220 pairs) to examine the heritability of the individual dimensions, their co-heritability and whether an independent or common pathway model fit the data best. If a common pathway model fit best, then shared etiological factors were mediated by a latent trait, while if an independent pathway model fit best, then shared etiological factors influenced dimensions directly. Determining which model fits best will help inform the optimal design for OCD genetic studies. We examined if removing the hoarding dimension affected the fit of common and independent models since the other OC dimensions may be a more cohesive etiological group (Mataix-Cols and Pertusa, 2012, Samuels et al., 2007). To be useful for genetic research, the TOCS should be at least as heritable as established measures of OC traits and, cluster into heritable OC trait dimensions similar to those previously reported (Bloch et al., 2008). If OC trait dimensions had both shared and unique genetic influences and were best explained by an independent model, this suggests research should not solely focus on a unitary OC trait, but also individual OC trait dimensions in youth.

## METHOD

### Sample & Research Design

We recruited 17,263 children and adolescents 6-18 years of age at the Ontario Science Centre, a local science museum in Toronto, Canada. Our final sample consisted of 16,718 participants with complete questionnaire information (mean age 11.1 years; SD +/− 2.8 years; 50.5% male). Informed consent, and verbal assent where applicable, approved by the Hospital for Sick Children Research Ethics Board were obtained from all participants. The authors assert that all procedures contributing to this work comply with the ethical standards of the relevant national and institutional committees on human experimentation and with the Helsinki Declaration of 1975, as revised in 2008. We collected behavioural information about participants (18.2%) if thought to be capable of self-reporting or from their parents (81.8%). Participants provided a saliva DNA sample using 2mL Oragene^®^ kits (DNA Genotek Inc, Ontario, Canada).

### Measures

A computerized, English questionnaire covered demographics, medical history, and OC traits measured by the TOCS and the Obsessive-Compulsive Scale of the Child Behavior Checklist (CBCL-OCS). The 21 TOCS items were scored on a scale of −3 to +3 (−3 far less often than average; −2 less often than average; −1 slightly less often than average; 0 average amount of time; 1 slightly more often than average; 2 more often than average; and 3 far more often than average). The TOCS has excellent internal consistency, inter-rated reliability, and divergent and convergent validity (Park et al., 2016). We created standardized TOCS z-scores for age and gender to generate a single standardized total score across informants. Total scores were modelled using linear regression controlling for age and gender, for parent- and self- respondents separately and residual scores were obtained. Participants were divided into 30 groups according to respondent (parent- or self-report), gender, and integer age groups. Parent respondent groups included integer ages from 6-15 and self-respondent groups included integer ages from ages 13-17. Standardized scores corresponding to the empirical percentile of each individual were assigned within each of the 30 groups separately. We also compared the heritability of the unitary OC trait construct measured in the TOCS to an established measure of OC traits: the CBCL-OCS (Nelson et al., 2001, Hudziak et al., 2006). The 8 items were each scored on a scale of 0 to 2 (0 not true; 1 somewhat/sometimes true; and 2 very/often true), and a total score was calculated by summing the item scores (range: 0-16).

### Twin Sub-Sample

We estimated heritability from 220 twin pairs. Their zygosity was initially determined by a twin questionnaire adapted from Cohen *et al.* (1975), and confirmed using a 16 marker microsatellite panel, following the protocol outlined in the study by Yang *et al.* (2006). DNA extracted from the saliva samples were analyzed for short tandem repeats using the AmpFLSTR^®^ Identifiler^TM^ PCR Amplification kit (PE Applied Biosystems, Foster City, CA, USA), a panel which consists of 15 autosomal, codominant, unlinked loci and the sex-determining marker, amelogenin amplified in a single PCR (Yang et al., 2006). Twin pairs were classified as monozygotic (MZ) if all 16 markers were identical between the pair; otherwise they were classified as dizygotic (DZ) (Yang et al., 2006). We had a total of four sets of DZ triplets. We randomly selected two siblings from each triplet to be a DZ twin pair and excluded the other member of the triplet. Our final twin sub-sample included 60 MZ twin pairs (50% male) and 160 DZ twin pairs (60 male, 33 female, and 67 opposite-sex pairs). The mean age of the twins was 10.5 years (SD +/− 2.6 years) and no individuals had a reported diagnosis of OCD.

### Statistical Analysis

#### Factor Analysis

Exploratory factor analysis with principal components using varimax rotation was conducted in SAS (version 9.3) to examine the underlying structure of the TOCS. We also conducted promax rotation because of the expected correlation of TOCS items. Pearson’s correlations were performed to examine the phenotypic correlations between the OC trait dimensions, using IBM SPSS Statistics 21.0 software.

#### Heritability Analyses

##### Univariate Models

Intraclass correlations for each trait and across traits within MZ and DZ twins were examined. The heritability of total OC traits and each individual OC trait dimension was estimated by structural equation modelling with age, sex, and respondent included as covariates using full information maximum likelihood (which included pairs with complete [n = 217] and incomplete data [n = 3]) in OpenMx (Boker et al., 2011). For analyses using standardized z-scores, age, sex, and respondent (parent or self) covariates were not included in the models because these factors were incorporated during z-scores calculation (see above). Saturated model fit was conducted to test the assumption of equality of means and variances between the MZ and DZ twins (Neale et al., 2006). The goodness of fit parameters used to compare twin models were the likelihood-ratio chi-square statistic (χ^2^) and Akaike’s information criterion (AIC).

We decomposed the total variance of the CBCL-OCS, TOCS total scores and each of the TOCS OC trait dimensions identified in our factor analysis into genetic and environmental factors. Genetic variance could be attributable to additive effects (A), and/or dominance (non-additive) effects (D). Environmental variance was partitioned to common environmental (C) influences, which are shared by family members, and unique environmental (E) factors, which also includes measurement error. In the ACE model, the within-pair genetic correlation was set at 1 for MZ twins and 0.5 for DZ twins. In the ADE model, the genetic correlation of MZ was still fixed at 1, but the genetic correlation of DZ twins was fixed at 0.25 (Plomin et al., 2012). The significance of the individual variance components was assessed by comparing the fit of the full models (ACE and ADE) to the nested sub-models (AE, CE, and E) where the effect was dropped.

To examine differences in heritability between the sexes, we observed intra-pair correlations by zygosity and sex. Our sample only had 33 DZ female twin pairs, and because the opposite-sex DZ twin correlations (0.40) were generally similar to the DZ same-sex twin correlations (0.47 for DZ males, 0.41 for DZ females) we did not further test sex differences in heritability. We could not examine differences in respondent (parent or self) because there were very few self-reporting twins (n = 54).

##### Multivariate Models

We tested the degree A, C, and E factors accounted for the covariance between the OC trait dimensions for the TOCS. We fit a multivariate correlated factor model to estimate the correlations between the A, C, and E variance components of the OC trait dimensions. A correlation between the A variance components of two measures was interpreted as an indication of a shared genetic basis, and a correlation between the C or E variance components was interpreted as an indication of overlapping environmental influences.

To understand how A, C, and E factors influence the co-variance between trait dimensions, we compared the correlated factor model to the common and independent pathway models (Kendler et al., 1987). In the common pathway model, the covariance of the OC trait dimensions is accounted for by a single latent phenotype influenced by shared additive genetic (Ac), common environment (Cc), and unique environment (Ec) factors. The model also estimates dimension-specific genetic (As), common environment (Cs), and unique environment factors (Es). The independent pathway model accounts for covariance of the dimensions by estimating a Ac, Cc, and Ec factor that directly influences each dimension (i.e., not through a latent phenotype) and dimension-specific variance is accounted by estimated As, Cs, and Es factors for each dimension. The best fitting model was selected using the AIC.

## RESULTS

### OC Trait Dimensions

With all 21 TOCS items, we initially obtained a 6 factor solution that included 14 items using varimax rotation where all primary factor loadings were >0.7 (factor with the highest loading) and no secondary loadings were >0.4 (all other factors). Two items, “experiences unwanted upsetting thoughts or images” (upsetting) and “spends time checking and rechecking homework” (homework) factored separately from the other items and were excluded from the final factor model. Upon re-examining the content and clinical relevance of these two items, the ‘upsetting’ item was too general, capturing a broad non-specific behavioural trait, and the ‘homework’ item was too specific although it was intended to capture a checking compulsion. The 14-item factor solution excluded items that queried common OC behaviours (e.g., counting) so we included 5 additional items to the 6 factor solution to include of as many of the items as possible. Our factor analysis of the 19 TOCS items resulted in six OC trait dimensions, which explained 75.7% of the variance: Cleaning/Contamination, Symmetry/Ordering, Superstition, Rumination, Counting/Checking, and Hoarding (Table 1). Descriptive statistics for these 6 factors can be found in Park et al. (2016). The results were similar for both parent- and self-report when examined separately (data not shown). The same factor structure was achieved using promax and varimax rotation.

**Table 1.** Factor analysis of the TOCS (19 items). The table shows factor loadings for each of the 19 items on the 6 OC dimensions.

| TOCS Item | Factor Loading |  |  |  |  |  |
| --- | --- | --- | --- | --- | --- | --- |
|  | Factor 1:<br>Cleaning/<br>Contamination | Factor 2:<br>Symmetry/<br>Ordering | Factor 3:<br>Superstition | Factor 4:<br>Rumination | Factor 5:<br>Counting/<br>Checking | Factor 6:<br>Hoarding |
| Wash | 0.84 | 0.09 | 0.09 | 0.13 | 0.19 | 0.06 |
| Germ | 0.84 | 0.17 | 0.18 | 0.10 | 0.09 | 0.10 |
| Clean | 0.80 | 0.13 | 0.17 | 0.15 | 0.17 | 0.05 |
| Dirt | 0.77 | 0.31 | 0.09 | 0.11 | 0.10 | 0.03 |
| Ruined | 0.49 | 0.40 | 0.37 | 0.01 | 0.07 | 0.25 |
| Interfere | 0.21 | 0.76 | 0.17 | 0.15 | 0.18 | 0.21 |
| Not Exactly | 0.28 | 0.66 | 0.19 | 0.42 | 0.11 | 0.16 |
| Symmetrical | 0.26 | 0.64 | 0.12 | 0.11 | 0.46 | 0.10 |
| Repeat | 0.24 | 0.60 | 0.29 | 0.24 | 0.35 | 0.13 |
| Bad Luck | 0.17 | 0.18 | 0.79 | 0.08 | 0.27 | 0.21 |
| Special | 0.14 | 0.25 | 0.74 | 0.11 | 0.34 | 0.05 |
| Healthy | 0.35 | 0.11 | 0.64 | 0.33 | 0.07 | 0.17 |
| Guilty | 0.17 | 0.19 | 0.14 | 0.82 | 0.25 | 0.11 |
| Thinking | 0.18 | 0.23 | 0.17 | 0.80 | 0.27 | 0.12 |
| Checks | 0.27 | 0.12 | 0.17 | 0.25 | 0.74 | 0.07 |
| Count | 0.17 | 0.40 | 0.26 | 0.15 | 0.66 | 0.15 |
| Do Certain | 0.09 | 0.20 | 0.26 | 0.22 | 0.71 | 0.15 |
| Throwing | 0.12 | 0.15 | 0.12 | 0.15 | 0.09 | 0.87 |
| Useless | 0.07 | 0.17 | 0.16 | 0.07 | 0.16 | 0.87 |

Phenotypic inter-factor correlations are shown in Table 2. The highest correlation was observed between the Counting/Checking and Symmetry/Ordering (*r* = 0.70). Hoarding showed low correlations with the other five dimensions (*r* = 0.31-0.52) and was less correlated with the TOCS total score than the other five dimensions (Table 2).

**Table 2.** Factor-Factor Phenotypic Correlations. Pearson’s correlation values between each of the six dimensions of the TOCS. The values on the left show correlations in the whole sample (N=16,718), and the bold values show correlations from the twin sub-sample (n = 220 pairs).

| TOCS Symptom Dimensions |  |  |  |  |  |  |  |  |  |  |  |
| --- | --- | --- | --- | --- | --- | --- | --- | --- | --- | --- | --- |
|  | Cleaning/<br>Contamination |  | Symmetry/<br>Ordering |  | Superstition |  | Rumination |  | Counting/<br>Checking |  | Hoarding |
| Cleaning/<br>Contamination | 1 |  | - |  | - |  | - |  | - |  | - |
| Symmetry/<br>Ordering | 0.62 | **0.59** | 1 |  | - |  | - |  | - |  | - |
| Superstition | 0.55 | **0.58** | 0.62 | **0.65** | 1 |  | - |  | - |  | - |
| Rumination | 0.44 | **0.45** | 0.60 | **0.64** | 0.51 | **0.59** | 1 |  | - |  | - |
| Counting/<br>Checking | 0.50 | **0.52** | 0.70 | **0.72** | 0.63 | **0.70** | 0.60 | **0.66** | 1 |  | - |
| Hoarding | 0.31 | **0.29** | 0.45 | **0.52** | 0.42 | **0.47** | 0.34 | **0.49** | 0.52 | **0.56** | 1 |
| TOCS Total<br>Score | 0.80 |  | 0.88 |  | 0.79 |  | 0.73 |  | 0.81 |  | 0.57 |
| p≤0.01 for all values |  |  |  |  |  |  |  |  |  |  |  |

### Univariate Heritability Models

Intraclass correlations for the TOCS total and dimension scores in the twins are shown in Table 3. No differences in the means and variances for the MZ and DZ twins were observed. For all variables, MZ twin correlations were larger than DZ correlations suggesting a genetic contribution to OC traits. MZ twin correlations were approximately two times large than DZ correlations across traits except for Cleaning/Contamination (Supplemental Table 1). Table 3 provides the standardized parameter estimates and the 95% confidence intervals (CI) for the ACE or AE models. The small sample size resulted in low power to detect small effects. For example, heritability of the Cleaning/Contamination dimension was 30%, which is not negligible, but was not found to be significant (95% CI: 0.0-0.7).

**Table 3.** Univariate Heritability Analyses of Overall OC Trait and its Dimensions. The table shows intraclass correlations (ICC) within the monozygotic (MZ) and dizygotic (DZ) twins followed by chi-square (χ^2^) differences, degrees of freedom (df) differences, and the p-values comparing the saturated model to the ACE model. A: additive genetic influence; C: common environmental influence; D: non-additive genetic (or dominance) influence; E: non-shared environmental influence. 95%-confidence intervals (CI) are shown. Δ = change in relevant statistic; AIC = Akaike’s information criterion.

| Variable | ICC | | Best Fitting model | $\Delta$ AIC | $\Delta \chi^2$ | $\Delta$ df | p-value | A (CI) | C (CI) | E (CI) |
| --- | --- | --- | --- | --- | --- | --- | --- | --- | --- | --- |
|  | MZ (CI)<br>(N=120) | DZ (CI)<br>(N=320) |  |  |  |  |  |  |  |  |
| TOCS |  |  |  |  |  |  |  |  |  |  |
| *Total Z-Score\** | 0.74<br>(0.60-0.83) | 0.37<br>(0.23-0.50) | AE | 7.48 | 6.52 | 7 | 0.48 | 0.74<br>(0.63, 0.82) | n/a | 0.26<br>(0.18, 0.37) |
| CBCL-OCS |  |  |  |  |  |  |  |  |  |  |
| *Total Score* | 0.62<br>(0.44-0.75) | 0.34<br>(0.20-0.47) | AE | 4.95 | 9.05 | 7 | 0.25 | 0.56<br>(0.40, 0.68) | n/a | 0.44<br>(0.32, 0.60) |
| TOCS Dimensions |  |  |  |  |  |  |  |  |  |  |
| *Cleaning/Contamination* | 0.56<br>(0.36-0.71) | 0.40<br>(0.27-0.53) | ACE | 7.60 | 4.40 | 6 | 0.62 | 0.30<br>(0, 0.66) | 0.26<br>(0, 0.52) | 0.45<br>(0.31, 0.63) |
| *Symmetry/Ordering* | 0.72<br>(0.57-0.82) | 0.30<br>(0.15-0.43) | AE | 8.67 | 5.33 | 7 | 0.62 | 0.70<br>(0.57, 0.79) | n/a | 0.30<br>(0.21, 0.43) |
| *Superstition* | 0.70<br>(0.54-0.81) | 0.46<br>(0.33-0.58) | ACE | 10.38 | 1.62 | 6 | 0.95 | 0.50<br>(0.17, 0.78) | 0.20<br>(0, 0.44) | 0.29<br>(0.20, 0.43) |
| *Rumination* | 0.55<br>(0.35-0.71) | 0.27<br>(0.12-0.41) | AE | 6.04 | 7.97 | 7 | 0.34 | 0.53<br>(0.36, 0.66) | n/a | 0.47<br>(0.34, 0.64) |
| *Counting/Checking* | 0.76<br>(0.62-0.85) | 0.37<br>(0.23-0.50) | AE | 7.22 | 6.78 | 7 | 0.45 | 0.77<br>(0.66, 0.84) | n/a | 0.23<br>(0.16, 0.34) |
| *Hoarding* | 0.66<br>(0.49-0.78) | 0.31<br>(0.16-0.44) | AE | 4.54 | 9.46 | 7 | 0.22 | 0.61<br>(0.46, 0.72) | n/a | 0.39<br>(0.28, 0.54) |

Heritability of OC traits was measured based on the TOCS z-score and the CBCL-OCS total score. Additive genetic factors accounted for 74% of the variance of OC traits measured by the TOCS z-score with 26% of the variance explained by unique environmental factors and measurement error. For the CBCL-OCS, the genetic contribution was 56%, with unique environmental factor and measurement error accounting for the remaining 44% of the variance.

As shown in Table 3, the AE model fit well for most of the dimensions, although the ACE model was more parsimonious than the AE model for the Cleaning/Contamination and Superstition dimensions based on lower AIC. Considerable genetic contributions were observed for all dimensions with heritability estimates ranging from 30-77%. For most of the dimensions, more than half of the variance was explained by genetic factors with the exception of Cleaning/Contamination, where approximately 70% of its variance was explained by environmental factors. For Cleaning/Contamination, 26% of the variance was explained by a common environmental effect.

### Multivariate Heritability Models

We examined the genetic correlations of the OC dimensions in our twin sample by decomposing the covariance between pairs of dimensions into genetic and environmental components to estimate the extent that these components influenced the dimensions. Genetic and environmental correlations between the dimensions from the multivariate analyses are shown in Table 4.

**Table 4.**
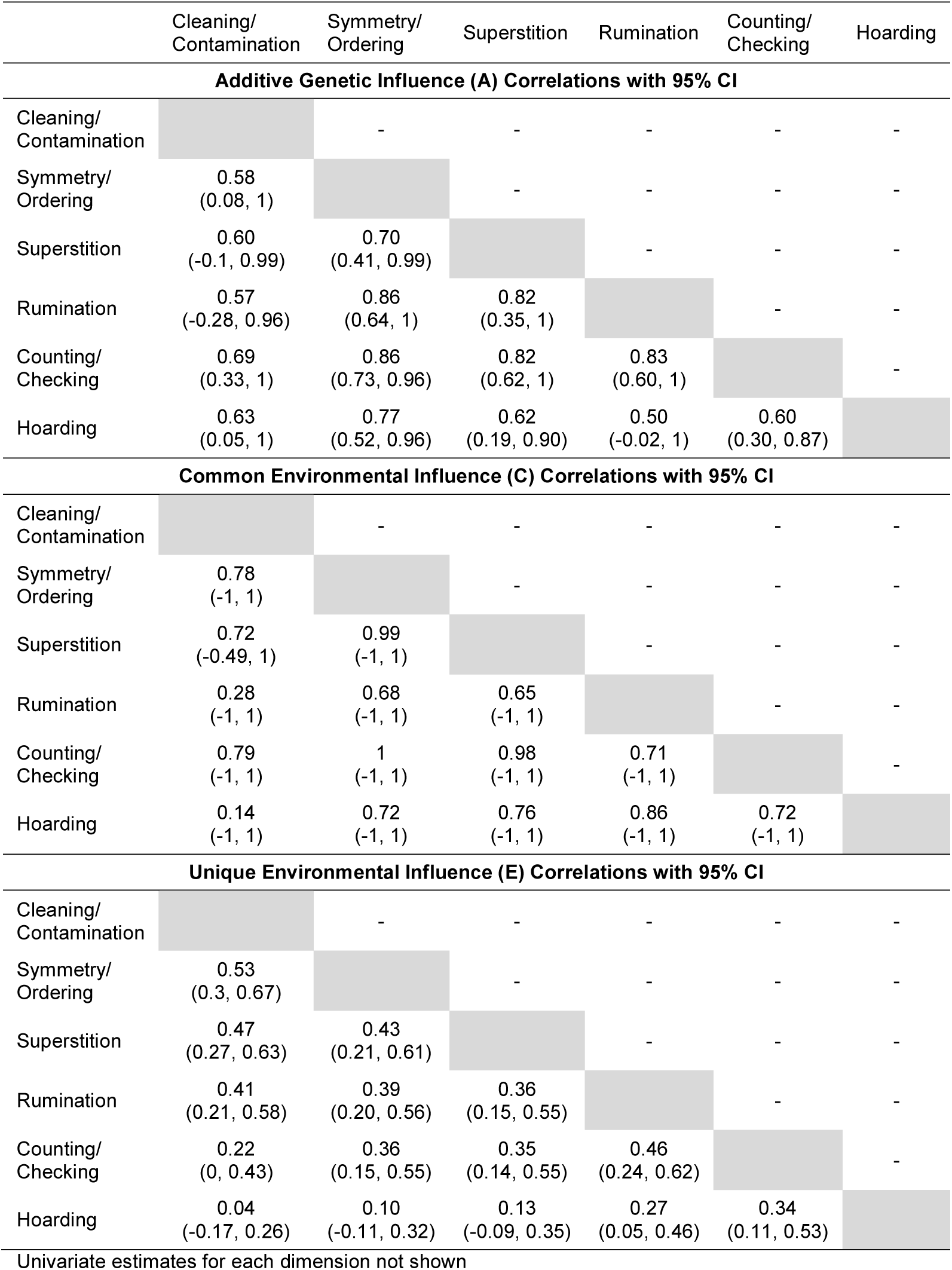
Multivariate Twin Analysis Matrices for all OC Dimensions. The table shows the correlations of additive genetic (A), common environmental (C) and unique environmental (E) variance between each of the obsessive-compulsive (OC) dimensions with 95%-confidence intervals (CI).

Additive genetic correlations between OC trait dimensions accounted for the majority of the co-variance of the dimensions. Significant correlations between A were observed for most pairs of dimensions except for Cleaning/Contamination with Superstition and Rumination. The highest additive genetic correlation was 0.86 observed for Symmetry/Ordering and Counting/Checking and for Symmetry/Ordering and Rumination dimensions. Unique environmental influences accounted for significant co-variance between OC trait dimensions as well. The Cleaning/Contamination and the Symmetry/Ordering dimensions showed highest E correlations (0.53). The lowest unique environmental correlation was seen for the Cleaning/Contamination and the Hoarding dimension (0.04).

We compared the fit of the ACE common pathway, independent pathway, and correlated factor models for the 6 OC trait dimensions. The independent pathway model fit best (AIC = 8243.88, df = 2595, *p* = 0.22) compared to the common pathway (AIC = 8259.76, df = 2604, *p* = 0.02) and correlated factor models (AIC = 8265.35, df = 2568). The model fitting was unchanged by removing the Hoarding dimension (data not shown). As shown in Figure 1, the majority of shared variance for each dimension was accounted for by genetic factors (Ac; 32-58%); except Cleaning/Contamination where common environment (Cc) accounted for the majority of the variance (36%). Genetic influences (As) accounted for the majority of dimension-specific variance for only Hoarding and Superstition (19-26%). For all other dimensions, unique environment (Es) significantly accounted for the majority of the dimension-specific variance (Es = 17-38%). Variance estimates with CIs from the independent model are presented in Supplemental Table 2.

**Figure 1:**
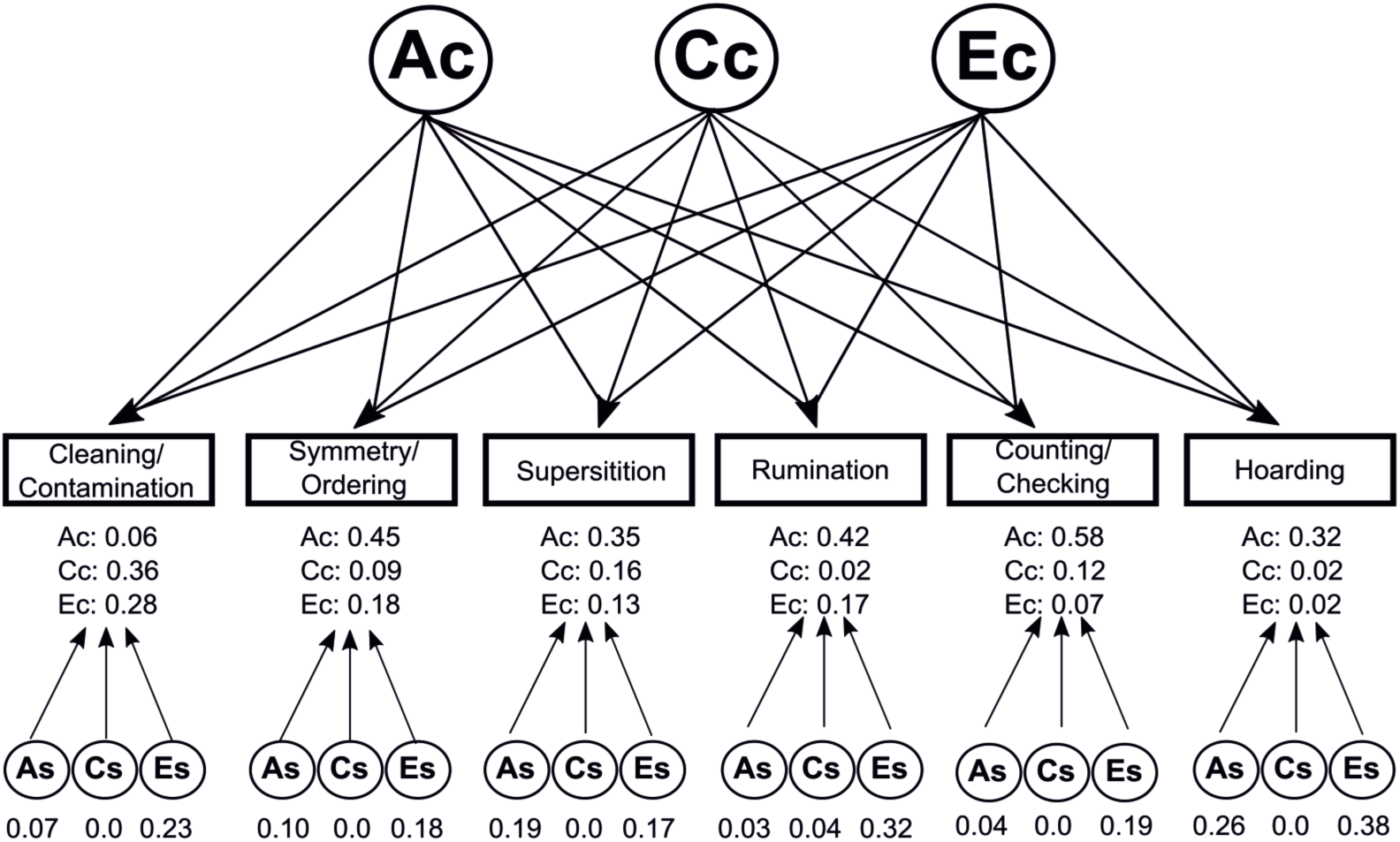
Independent Pathway Model of OC Dimensions. The best fitting model of obsessive-compulsive (OC) dimensions was the independent pathway model where shared co-variance was mostly attributed to shared additive genetic influences (Ac) and to a lesser degree by unique environment (Ec). Dimension-specific variance was most explained by unique environment (Es) and, to a lesser degree, additive genetic factors (As) for Hoarding and Superstition. Shared (Cc) and dimension-specific (Cs) common environment did not explain much variance except for Cleaning/Contamination.

We also compared the heritability of the 6 dimensions with the initial 6 factor structure using 14 items and the 6 factor structure using 19 item factor that included more clinically-relevant items. Univariate estimates of heritability were similar for the two factor solutions. An exception was for Symmetry/Ordering which best fit an ADE model rather than an AE model in the 19-item solution (Supplemental Table 3). Genetic correlations between dimensions were also similar regardless of whether 14 or 19 items were included (Supplemental Table 4). The independent model still fit best (AIC = 7258.44, df = 2595, *p* = 0.39) compared to the common pathway (AIC = 7259.80, df = 2604, *p* = 0.09) and correlated factor models (AIC = 7284.12, df = 2568). The proportion of A, C, E for shared and dimension-specific variance was similar to the 19-item factor model (Supplemental Table 5). For Cleaning/Contamination, genetic influences accounted for most of the shared variance while unique environment accounted for most of the dimension-specific variance. For Superstition, unique environment accounted for most, while genetic influences accounted for none, of the dimension-specific variance.

## DISCUSSION

One strategy for improving the power of genetic studies in OCD is to focus on OC traits, which are widely distributed in the general population, rather than limit samples to participants with a clinical OCD diagnosis. We developed the TOCS to measure the full range of OC traits in children and adolescents (Park et al., 2016). To be suitable for genetic research of OCD, the TOCS should capture accepted OCD dimensions and should be heritable. The TOCS factored into 6 heritable and co-heritable OC dimensions similar to those reported from studies using traditional OCD scales (Bloch et al., 2008, Stewart et al., 2008). We further used the TOCS to explore whether OC dimensions in youth were accounted for by a latent trait and the degree to which these dimensions shared etiological factors using ACE twin models comparing independent pathway and common pathway models. We showed that OC dimensions had both shared and unique genetic influences that were best explained by an independent model suggesting that the shared genetic influences on OC dimensions were not mediated by a latent trait. Hoarding was phenotypically distinct from the other dimensions but was still genetically correlated and shared part of its additive genetic influences with other OC dimensions. Thus, although hoarding was somewhat distinct, a common underlying etiology was present. Together our results show that the TOCS is a useful tool for genetic OCD research and because OC dimensions have common and distinct underlying etiologies, they should be studied together and individually in genetic research.

Our new TOCS measure captured common OCD dimensions. Most paralleled the dimensions identified in previous factor analysis studies (Bloch et al., 2008, Stewart et al., 2008, Stewart et al., 2007, Ivarsson and Valderhaug, 2006, McKay et al., 2006) although we identified separate Counting/Checking and Rumination dimensions. Counting and Checking symptoms often cluster with other symptom types including Symmetry/Ordering and/or Hoarding (Bloch et al., 2008). The Rumination dimension contained items consistent with those of the sexual/religious and obsessions dimensions from previous studies (Moore et al., 2010, Ivarsson and Valderhaug, 2006, Mataix-Cols et al., 2008). Overall, there was considerable convergence in the factor structure of the TOCS and traditional OCD scales.

The TOCS also showed similar heritability to previous OCD measures. The estimated heritability of the TOCS total score was 74%, which is higher than estimates for OC traits from previous twin studies (Bolton et al., 2007, Hudziak et al., 2004, van Grootheest et al., 2008a, van Grootheest et al., 2005, Jonnal et al., 2000, Van Grootheest et al., 2007b). The TOCS was as heritable as the CBCL-OCS, an established heritable OC trait measure (Hudziak et al., 2004), supporting the utility of the TOCS for genetic research.

All OC trait dimensions were also heritable, similar to previous studies in youth and adults (Moore et al., 2010, Iervolino et al., 2011, Katerberg et al., 2010, van Grootheest et al., 2008b, Jonnal et al., 2000, Mathews et al., 2007). Environmental factors only significantly contributed to the Cleaning/Contamination dimension suggesting a distinct etiological mechanism. Low additive genetic effects were reported previously for this dimension in an adolescent population-based sample (Moore et al., 2010). The effect of common environment may result from family values, education or parental modeling.

Phenotypic heterogeneity in the TOCS, demonstrated by 6 OC dimensions, also reflected some etiological heterogeneity. Genetic factors contributed considerably to all OC dimensions, although less so for Cleaning/Contamination. All OC dimensions were also co-heritable indicating that they share some genetic influences. However, how much genetic influences were shared between dimensions and affected dimensions individually varied. For many OC dimensions, genetic influences were accounted for by shared effects with the other dimensions suggesting that similar genetic factors play an important role across phenotypically separate OC dimensions. In contrast, what made the OC dimensions different was accounted for mostly by non-shared environment rather than genetic factors. A notable exception was Hoarding, which showed considerable genetic effects that were shared with the other dimensions but also had considerable hoarding-specific genetic influences. A similar pattern was observed for Superstition. In a previous study of female adults (Iervolino et al., 2011) genetic effects contributed to more dimension-specific variance than in the present study although in both studies, unique environment accounted for the most variance specific to each dimension. One implication is that genetics may play a larger role in what makes dimensions similar while unique environment may play a bigger role in what makes dimensions different. Another is that the proportion of genetic and environmental factors differs across dimensions.

Additional evidence that OC dimensions have similar but separate etiological influences is our finding that the dimensions were best accounted for by an independent pathway model. If a common pathway model fit best, shared etiology of the OC dimensions would have been attributable to a latent trait (e.g., OC traits) suggesting that latent trait may be more useful in genetic studies than individual dimensions. Results from previous studies in adults on the fit of the common and independent pathway for OC traits are mixed (Iervolino et al., 2011, van Grootheest et al., 2008b). An important difference seems to be the number of OC dimensions captured by the scale they used – an independent model fits best when there was a broader spectrum of dimensions (Iervolino et al., 2011) while a common model fit best when there were fewer dimensions (van Grootheest et al., 2008b). Our finding that shared etiological factors contribute to OC dimensions in youth without being mediated by a latent trait suggests that simply measuring overall OC traits will not uncover the full spectrum of genetic influences on OC dimensions.

Hoarding is a symptom of OCD but also considered distinct in the DSM-5 (American Psychiatric Association, 2013). Our study supported the phenotypic distinction of hoarding but not a complete etiological distinction. Hoarding did not correlate well with the other dimensions or the TOCS total score but was genetically correlated with all other OC trait dimensions. Hoarding also had a distinct etiological profile with genetic influences accounting significantly to both shared and dimension-specific variance. Excluding Hoarding from the independent pathway model did not affect model fit suggesting that hoarding was not obscuring a latent OC trait that accounted for the other OC dimensions. In a previous adult twin study, hoarding and total OC symptoms were not highly genetically correlated but did share additive genetic effects (Mathews et al., 2014). Classifying hoarding as a distinct condition may be useful in the clinic and to find hoarding-specific mechanisms, but results indicate that hoarding likely shares considerable genetic risk with other OC trait dimensions and OCD and thus should be considered in OC genetic studies. Disorders may share their genetic etiology even when phenotypically and clinically distinct (Cross-Disorder Group of the Psychiatric Genomics, 2013).

Our factor analyses clearly demonstrated 6 OC dimensions, however, the number of items to include was less clear. Although the 14-item model fit best, it excluded items that represent common OC symptoms such as counting. Regardless of the number of items included, our genetic results were similar for the heritability and co-heritability of the dimensions in that genetic effects accounted for considerable variance for each dimension. The independent model fit the data best and most of the shared and dimension-specific variance were similar. One notable difference was that with fewer items, genetic rather than common environment accounted for more of the shared variance for Cleaning/Contamination. This suggests that the relative contribution of genetic and common environmental effects is less clear for Cleaning/Contamination, a conclusion that is supported by our univariate models and previous literature (Moore et al., 2010, Iervolino et al., 2011). The fact that almost all of the heritability results were similar with either factor structure suggests that the 6 dimensions are robust and heritable.

Our results should be examined with the following considerations. Despite a relatively small sample of twins, we found statistically significant heritability and co-heritability of OC traits and trait dimensions. However, power for detecting sex or age differences was limited (Moore et al., 2010, Hudziak et al., 2004, van Grootheest et al., 2008a).

The TOCS is useful for studying OC traits. This measure identified several heritable OC dimensions similar to those in previous studies. Our results support the hypothesis that OC traits are phenotypically and etiologically heterogeneous. OC dimensions that co-occur can have different etiological mechanisms and those less well correlated phenotypically, like hoarding and other dimensions, could nevertheless share genetic risk. To uncover the genetics of OC traits and OCD, OC trait dimensions should be considered both individually and together.

## Acknowledgements

This work was part of LSP’s Master’s thesis at the University of Toronto. We thank Lisa Strug for statistical consultation and all individuals involved in collecting this dataset.

